# N-Gene and T-Gene Deregulation Networks: A data-driven causal framework for the analysis of gene interventions in cancer

**DOI:** 10.1101/2023.12.28.573550

**Authors:** Frank Castel, Roberto Herrero, Jean Pierre Gómez, Gabriel Gil, Augusto Gonzalez

## Abstract

**Background:** The construction of realistic gene regulatory networks is currently hindered by the complex, often redundant, combinatorial and multi-layered nature of gene interdependencies, the limited availability of functional annotations, and the intrinsic shortcomings of co-expression-based approaches. Here, we introduce Gene Deregulation Networks (GDNs), a new structure in which linked genes indicate that an expression deregulation in one is likely to propagate to the other. GDNs are inferred directly from gene expression data using a probabilistic theory of causation, without requiring prior functional annotation or other domain knowledge.

**Methods:** Using highly specific tissue markers—N-genes specifically expressed in normal tissues and T-genes specifically expressed in tumors—we construct separate GDNs for normal and tumor tissues. Data are obtained from TCGA RNA-Seq profiles of bulk tumor and normal tissue samples across multiple cancer localizations. GDN links are preliminarily identified via a statistical test of causal sufficiency between gene expression deregulation events, and then reassessed as spurious or redundant using additional causal criteria. A simple scheme based on the GDNs is implemented in order to describe the evolution dynamics.

**Results:** The T-GDN in prostate adenocarcinoma comprises 6138 genes and 102362 directed edges (0.27% of possible connections). Genes with low expression deregulation frequency (< 0.2) exhibit high out-degrees, indicating high deregulation potential, while high-frequency genes (> 0.4) show high in-degrees, suggesting they are convergence points of deregulation cascades. *EPHA10* (frequency 0.74, in-degree 182) and *ENSG00000275479* (out-degree 209) represent these extremes. The N-GDN contains 1097 genes and 4984 edges, featuring compressed nodes reflecting the multi-step nature of somatic evolution. Analysis of tumor samples reveals that early-stage tumors rely primarily on spontaneous gene activations, while advanced tumors exhibit extensive cascade-driven deregulation. Preliminary simulations show qualitative agreement with experiments on gene knockdown in tumor cellular lines.

**Conclusions:** GDNs provide a robust framework to understand cancer progression through deregulation cascades. The separation into N and T networks, connected by NT-genes, i.e. genes common to both networks, offers a systematic basis for modeling carcinogenesis and the possible outcomes of therapeutic interventions.

## 1. Introduction

Current attempts to construct gene regulatory networks (GRNs) typically involve dozens of genes and a limited number of interactions among them [1,2]. However, experimental evidence demonstrates that perturbations affecting a few genes can trigger extensive signaling pathways and restructure the expression profile of tumors. For instance, the combined application of two interferons on a glioblastoma cell line produces widespread transcriptional changes [3], suggesting that realistic GRNs encompass many genes with numerous interconnections.

In this article, we introduce Gene Deregulation Networks (GDNs), inferred from expression data using the probabilistic theory of causation [4,5,6]. The underlying concept differs from traditional GRNs: rather than seeking functional relationships between genes or co-varying gene expression patterns, we address a simple question: does the inferred probability that a gene E is deregulated increase with the concurrent deregulation of another gene C? If so, we interpret this as a causal influence from C to E and connect the two genes within the GDN. Such inferences must ultimately be biologically validated. However, the fact that substantial fractions of known genes lack functional annotations justifies a fully data-driven, non-parametric approach to the GDN construction. Moreover, representing gene expression in terms of deregulation events rather than continuous values allows for a more biologically grounded account of gene-specific transcriptional signals and their downstream effects, without requiring prior functional annotations or domain knowledge.

We consider all genes (approximately 60,000 in The Cancer Genome Atlas [TCGA] data [7]) and assess possible causal relationships within all ordered gene pairs. An explicit construction of such networks for prostate adenocarcinoma (PRAD), excluding non-causal statistical correlations, was performed in [8]. However, in that initial attempt, deregulation was defined in terms of differential expression, thereby introducing artifacts.

In Refs. [9,10], N-genes and T-genes were introduced as expression markers for normal tissues and tumors, respectively. These markers are not based on differential expression but on a broader concept of expression deregulation. Typically, any tissue contains approximately 1,000 N-markers and 10,000 T-markers. About 500 genes are common to both groups, designated NT-genes. The remaining 49,000 genes do not distinguish normal from tumor samples, including, although not limiting to, genes that are not expressed in the tissue.

In Ref. [10], it was argued that somatic evolution of a normal tissue is taken to involve only N-genes, while clonal evolution of tumors should involve only T-genes. Thus, we can use expression data from normal samples to infer the N-GDN and tumor samples to infer the T-GDN. This approach is reasonable because N and T attractors are well-separated in gene expression space [11,12]. Only during normal-to-tumor transitions or forced tumor-to-normal phenotype reversal processes both networks participate, with NT-genes acting as effective bridges between them.

In the present paper, we construct and characterize N- and T-GDNs for multiple tumor localizations using TCGA data. The paper is organized as follows. In the Methods section, we summarize the definitions of N-, T- and NT-genes, and introduce GDNs within a causal discovery framework. We also describe the datasets and computational procedures used for constructing and analyzing the networks. Next, we present results for N- and T-GDNs across multiple cancer types and tissue localizations, emphasizing the occurrence of deregulation cascades in T-genes. Finally, we discuss the carcinogenic dynamics within the obtained networks and offer concluding remarks on the potential implications of our results for gene-level therapeutic interventions and for improving our understanding of cancer.

## 2. Methods

### 2.1 Tissue Markers from Gene Expression Data

Following [9,10], we define N-genes, T-genes, and NT-genes based on their expression patterns across normal and tumor samples. Genes not classified within these groups are not included in the present framework.

For each gene, expression values are mapped to discrete expression states with values e = -1 if expression is specific to normal samples (N-active), e = 1 if specific to tumor samples (T-active), and otherwise e = 0 (inactive). A Fisher test is then applied to assess whether the observed frequencies of N-active and T-active instances are statistically significant for each gene [9]. We use significance thresholds of 5% of total normal samples for N-activity and 10% of total tumor samples for T-activity [10].

Genes with statistically significant N- or T-activity are classified as N-genes or T-genes, respectively. NT-genes are those exhibiting both significant N- and T-activity across samples. Pure N- or T-genes exhibit significant activity in only one condition and are therefore active only in their corresponding state or inactive otherwise. Notably, an expression deregulation of a T-gene coincide with a T-active state (e = 1); however, in the case of N-genes, it correspond to an inactive state (e = 0) or, in the case of NT-genes, also a T-active state (e = 1).

### 2.2 Gene Deregulation Networks

T-genes and N-genes are nodes of the T- and N-GDNs, respectively. The construction of both networks follows a similar scheme. First, we build up a gene-by-sample matrix with N-genes and normal samples in the case of N-GDNs, and T-genes and tumor samples in the case of T-GDNs. Each entry of the matrix is assigned a binary indicator of whether an expression deregulation occurs or not [see last section for the definitions of N- and T-gene deregulation based on discrete expression states]. Genes with identical deregulation profiles (hereafter referred to as block genes) are indistinguishable within our framework and are therefore merged into a single representative gene. Then, we compute the conditional and unconditional frequencies of deregulation across samples. The frequency of deregulation for a given gene i, denoted v_i, is simply the ratio of the number of deregulations to the total number of samples in the matrix. The *conditional* frequency of a deregulation at gene j *given* a deregulation at gene i, denoted v_j|i_, is computed as v_ji_ / v_i_, where v_ji_ is the frequency of co-occurring deregulations at i and j. With these notions in place, a probabilistic causal sufficiency relation between genes i and j exists if and only if:

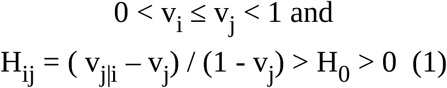

where H_ij_ is known as the Loevinger coefficient [13,14] and H_0_ is a positive real number between 0 and 1, independent of the pair of genes. If conditions (1) are satisfied, an arc i → j is added to the GDN.

Notice that H_ij_ > 0 entails v_j|i_ – v_j_ > 0, which is the main postulate of the probabilistic theory of causation: causes increase the probability of their effects [4,5,6]. Unlike v_j|i_ – v_j_, H_ij_ is largely independent of the frequency of putative effects and therefore better suited as a figure of merit for causation. We impose the more stringent condition H_ij_ > H_0_ > 0 to remove statistically insignificant instances of causal correlation. Our causal sufficiency criterion captures the inherent asymmetry of cause and effect in the prototypical case of v_i < v_j, since H_ij_ > H_ji_, thereby making inference from a deregulation at i to a deregulation at j more justified than the reverse. In this sense, we refer to causal sufficiency (a deregulation at i is sufficient but not necessary to observe a deregulation at j). Of course, the inference is probabilistic and allows for errors (instances where i is deregulated but j is not, despite H_ij_ > H_0_). In our computational protocol, we choose H_0_ = 1/2, which ensures that the effect occurs more than half the time when the cause is present. For a given pair of genes (causally correlated or not), H_ij_ can be higher or lower than H_0_, and even negative, which makes the criterion in (1) operationally meaningful. However, H_ij_ ≤ 1, with H_ij_ = 1 corresponding to the case where no inferential errors take place: any deregulation at i is accompanied by a deregulation at j. Taken together, this discussion supports the interpretation of H_ij_ > H_0_ as signaling a *prima facie* causal sufficiency relation between the involved genes [8].

When the putative effect, i.e., the deregulation at gene j, always takes place, ascribing causal sufficiency relations to it becomes meaningless (anything can cause its deregulation), hence the condition v_j_ < 1 in (1), which excludes such cases. Likewise, a putative cause must take place at least once (v_i_ > 0). In practice, this means that genes with frequency v_i_ = 0 or v_i_ = 1 are represented by isolated nodes in the GDN. The minimum frequency of expression deregulation for T-genes is 0.1, but v_i_ = 1 is still possible. The situation for N-genes is the opposite since our significance threshold imposes an upper but not lower bound on the frequency of expression deregulation, namely, we may have v_i_ = 0 but not v_i_ = 1.

To briefly illustrate the implications of our choice H_0_ = 1/2, we consider the Fisher exact test associated with contingency tables satisfying conditions (1). Specifically, we evaluate the probability (p-value) that, for two T-genes at the minimum frequency threshold in PRAD data (∼500 tumors, hence ∼50 deregulations per gene), a contingency table meeting these conditions arises by chance. The resulting p-value is on the order of 10_-18_. In contrast, setting H_0_ = 1/8 yields a p-value of 0.15. These results indicate that pairs of T-genes satisfying our prima facie causal sufficiency criterion for H_0_ = 1/2 are highly unlikely to arise from random fluctuations.

Some causal links (i → j) discovered by the previous conditions may be spurious, arising from non-causal statistical correlations between effects of a common cause k (i.e., k → i and k → j) [5]. A Reichenbach-like test helps identify and remove such cases [8]. Other links may be redundant due to causal transitivity if i → j is explained by the presence of i → k and k → j. We prune these by appealing to the Mokken test of double monotony [14]. The connection between the Mokken test and causal transitivity is discussed in Refs. [8,14a]. Additionally, a heuristic test, originally suggested by Mokken as well [14], is applied to define the orientation of links in cases where v_i_ = v_j_ and H_ij_ = H_ji_ [8,14a]. The application of these procedures typically yields a sparse, directed acyclic graph. Since H_ij_ > H_0_ implies H_ji_ > H_0_ when v_i_ = v_j_, we emphasize that the directedness and acyclicity of the resulting graph arise from running the full causal discovery algorithm (including arc-addition and arc-removal stages) on empirical data. Indeed, although the Loevinger coefficient, when applied to genes with distinct frequencies, guarantees the arc direction, causal asymmetry in the case of equal frequencies may or may not be recovered after the arc-removal stage, depending on the data. Moreover, directedness, acyclicity, and sparsity are strongly dependent upon the arc-removal stage.

### 2.3 Data, Software and Post-processing

TCGA level 3 expression data were obtained for PRAD [16], head and neck squamous carcinoma (HNSC) [17], lung adenocarcinoma (LUAD) [18], uterine corpus endometrial carcinoma (UCEC) [19], and lung squamous cell carcinoma (LUSC) [20] through the Genomic Data Commons portal. These datasets contain RNA-Seq expression profiles with ∼60,000 genes for normal and tumor tissue samples. The number of normal and tumor samples is 52 and 499 for PRAD, 44 and 502 for HNSC, 59 and 535 for LUAD, 23 and 552 for UCEC, and 49 and 502 for LUSC, respectively. Normal samples are derived from cancer patients. In some cases, patients contribute both normal and tumor samples to the dataset.

For the construction of GDNs, we rely on an original in-house C++ software called, CChains [8,15]. CChains operates on a binary matrix, whose entries, in our case, represent expression deregulations across genes and samples, and outputs the adjacency lists of the resulting graph.

Further network post-processing is carried out leveraging the established graph analysis engine NetworkX [21]. Using this library, GDNs are characterized by global and local network metrics, such as the distribution of node in- and out-degrees, the network diameter, the number of isolated, orphan, and child-less nodes, among others. In- and out-degrees correspond to incoming and outgoing causal connections, respectively. Furthermore, deregulation cascade analysis requires identifying, for each sample, genes that are deregulated without concurrent deregulation of their causal parent nodes within the network. We refer to these as spontaneously activated genes.

## 3. Results & Discussion

### 3.1 The T-Gene Deregulation Network in PRAD

Using TCGA-PRAD data [16], we constructed a T-GDN with 6138 nodes, 151 of which are isolated. The resulting network contains 102362 directed edges, representing a sparse network with edge density of 0.27%.

Figure 1 shows a scatter plot of node degrees vs. activation frequencies. For example, the *EPHA10* gene exhibits the highest deregulation frequency (0.74) and a very high in-degree (182), and a gene in the low-frequency region (namely, *ENSG00000275479*) shows the highest out-degree (209). Nodes with high out-degrees are typically found in the v < 0.2 region. In particular, we find 733 orphan nodes (in-degree = 0). Low-frequency genes thus have high deregulation potential, being capable of influencing many downstream genes. Conversely, high-frequency genes exhibit high in-degrees, suggesting their possible role as gene hubs in the GDN. There are 1775 child-less nodes (out-degree = 0).

**Figure 1:**
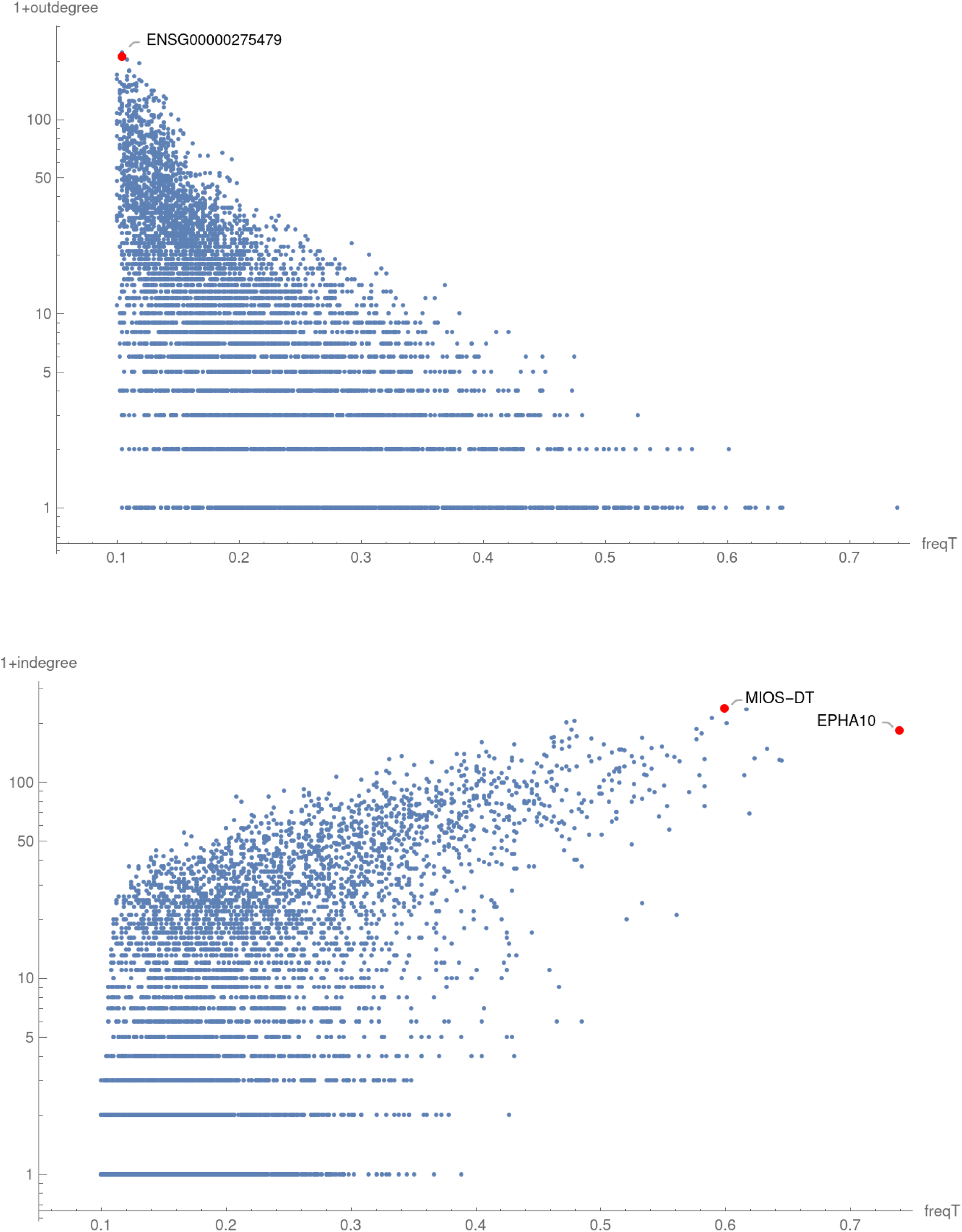
Characterization of the T-GDN in PRAD. Each point represents a gene. Node in-degrees and out-degrees are plotted against frequencies. EPHA10 shows the highest activation frequency (0.74) and a high in-degree (182). MIOS-DT exhibits the highest in-degree, 237. A gene in the low-frequency region, ENSG00000275479, shows the highest out-degrees (209).

Table I presents global and local network metrics. Supplementary Table I contains node properties (frequencies, in/out-degrees, gene identities). Supplementary Table II details causal relationships, indicating children nodes for each parent; parent nodes of a given gene can be obtained by transposition.

**Table I:** Some global and local metrics in the T-GDN and N-GDN in PRAD, HNSC, LUAD, UCEC y LUSC. The same values of minimal frequency, 0.1 and 0.05, are used to define T- and N-genes, respectively, in all the localizations.

|  | PRAD | HNSC | LUAD | UCEC | LUSC |
| --- | --- | --- | --- | --- | --- |
| **T-GDN** |  |  |  |  |  |
| Number of T-genes | 6138 | 9340 | 17037 | 16469 | 18596 |
| Sparsity | 0.0027 | 0.0022 | 0.0019 | 0.0018 | 0.0029 |
| Max freq_T | 0.74 | 0.89 | 0.98 | 1 | 1 |
| Mean diameter | 1.9 | 2.5 | 3 | 3.2 | 5.1 |
| Isolated nodes | 151 | 193 | 28 | 11 | 1 |
| Mean in-deg = Mean out-deg | 16.6 | 20.9 | 32.1 | 29.6 | 53.8 |
| Max in-deg | 237 | 400 | 1008 | 1758 | 1178 |
| Max out-deg | 221 | 160 | 300 | 341 | 426 |
| Child-less nodes (out-deg = 0) | 733 | 802 | 394 | 349 | 40 |
| Orphan nodes (in-deg = 0) | 1775 | 3103 | 4949 | 5281 | 5603 |
| **N-GDN** |  |  |  |  |  |
| Number of N-genes | 1097 | 1372 | 492 | 1192 | 1269 |
| Rank (number of nodes) | 499 | 774 | 470 | 773 | 1228 |
| Sparsity | 0.0200 | 0.0139 | 0.0151 | 0.0116 | 0.0093 |
| Max number of genes in a block node | 212 | 93 | 4 | 131 | 11 |
| Max freq_N | 0.31 | 0.5 | 0.81 | 1 | 0.9 |
| Mean diameter | 2.2 | 2.8 | 2.3 | 4.8 | 4 |
| Isolated nodes | 1 | 0 | 0 | 1 | 0 |
| Mean in-deg = Mean out-deg | 10.0 | 10.7 | 7.1 | 9.0 | 11.4 |
| Max in-deg | 60 | 56 | 27 | 31 | 40 |
| Max out-deg | 165 | 153 | 39 | 63 | 61 |
| Child-less nodes (out-deg = 0) | 93 | 121 | 81 | 87 | 136 |
| Orphan nodes (in-deg = 0) | 10 | 10 | 6 | 5 | 9 |

### 3.2 Deregulation Cascades Apparent in Tumor Samples

In this section, we map the deregulation profile of each sample onto the established T-GDN. The purpose of this mapping is to investigate how deregulations are distributed within the network on a sample-by-sample basis. As a result, we find that more than 90% of samples contain causally connected deregulations, which we term deregulation cascades.

To illustrate this, Figure 2 shows the total number of deregulated T-genes in each sample against the number of spontaneously activated genes in that sample—i.e. nodes that are deregulated within the sample, without deregulation in their causal parents. By definition, spontaneously activated genes can only be activated by factors external to the T-GDN.

**Figure 2:**
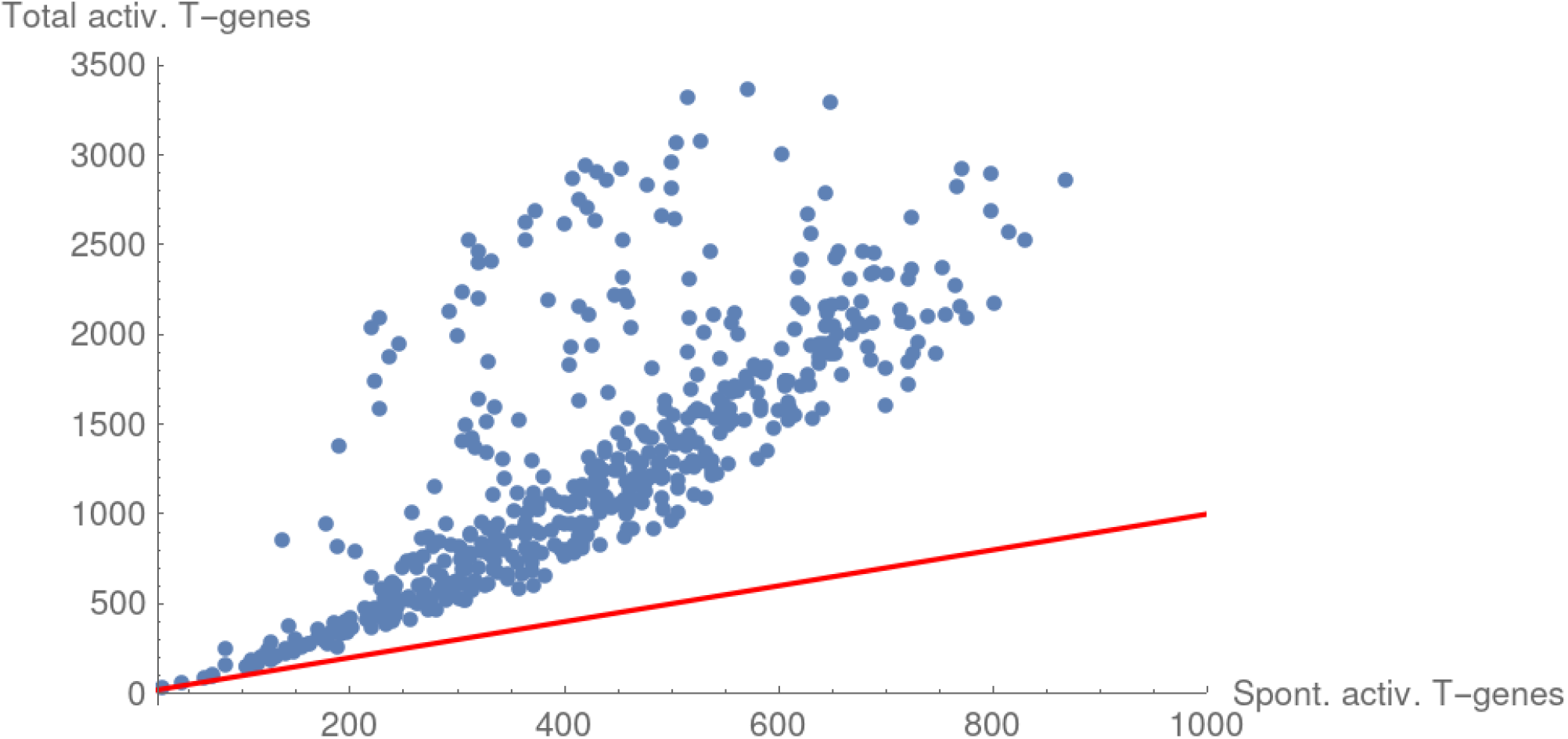
Total number of activated T-genes in samples versus number of spontaneously activated genes. (nodes presumably not deregulated by cascades). Each point represents a sample. The diagonal (red) provides visual reference. Early-stage tumors (lower left) show predominance of spontaneous activation; advanced tumors (upper right) show increasing cascade contributions.

The diagonal in Figure 2 provides a reference for the importance of spontaneously activated genes. When the deregulated nodes are few in number—a feature of newly formed tumors [10]—most deregulations are spontaneous. As tumors evolve, more nodes are deregulated [10]. Such an increase in the number of deregulations is associated with a departure from the diagonal in Figure 2, indicating that at least some cascades extend beyond their onsets. The latter result suggests deregulation cascades as a prominent mechanism behind tumor progression.

To survey the transition from a normal tissue to a tumor, we rely on paired normal and tumor samples from the same patient within PRAD data. For example, Figure 3 shows a representation of the deregulation profiles of one such pair of normal and a tumor samples. In the top panel, each point corresponds to a deregulated (or activated) T-gene, whereas in the bottom panel, each point corresponds to an active N-gene (i.e., an N-gene not exhibiting deregulation in that sample). The y-axis indicates the out-degree of the N- and T-genes in their respective GDN. The frequency of T-gene deregulations and the frequency of N-gene deregulations are shown on the x-axis of the top and bottom panels, respectively.

**Figure 3:**
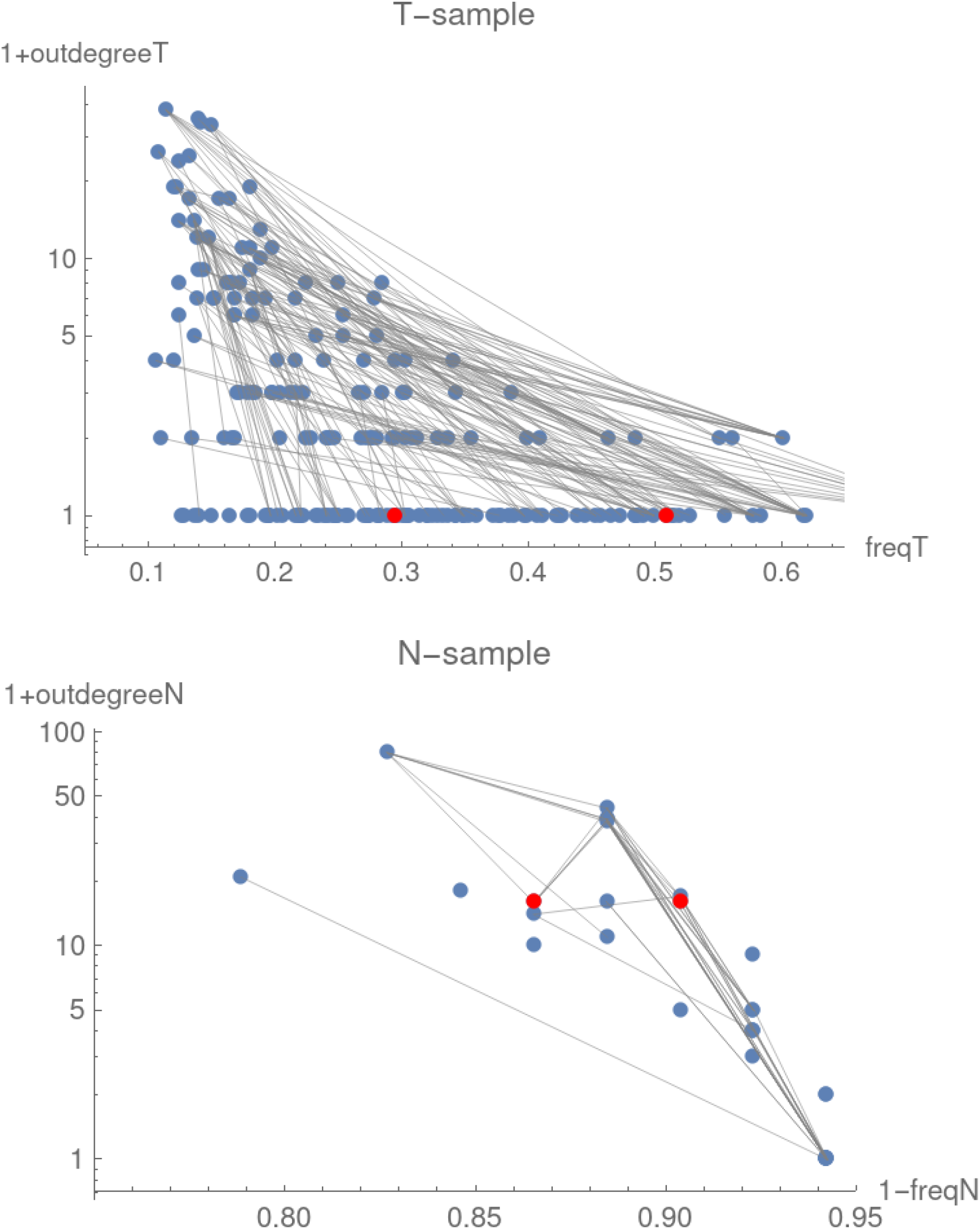
Activated nodes in PRAD paired samples. **Top panel:** Activated nodes in the tumor sample, showing deregulation cascades (paths) and spontaneously activated genes. **Bottom panel:** Activated N-nodes in the corresponding normal sample. Deactivation pressure from already-deactivated nodes (not shown) also affects activated N-genes. Common NT-genes are highlighted in red.

In the tumor sample, there are 201 deregulated (i.e., activated) T-genes, revealing its relatively early stage. Two of these 201 are NT-genes which are N-active in the normal sample. We marked them in red in both panels. These shared nodes between the N- and T-GDNs are consistent with a switch from N to T activity during the normal-to-tumor transition [10].

Figure 3 also provides a partial representation of the T- and N-GDN. In particular, we show the causal links among deregulated T-genes and non-deregulated N-genes in the top and bottom panels of Figure 3, respectively. Both networks are highly structured although the activated T-GDN seems more complex, even if still sparse (see Table I). In particular, the top panel of Figure 3 shows that deregulation cascades do not form simple linear causal chains but instead are organized as complex branching structures.

Recall that each tumor sample can be classified by the activation status of a complete 8 T-gene panel [9,10]. This earlier result can be reinterpreted as follows. Although deregulations in a tumor sample can be large in number (30 - 3300) and distributed according to intricate patterns within the T-GDN (as apparent in Figure 3), tumor heterogeneity can be summarized by the activation state of only 8 genes [10]. This suggests that cancer-relevant deregulation cascades within each activation pattern of the T-GDN necessarily intersect at least one of these 8 genes.

### 3.3 T-Gene Deregulation Networks in HNSC, LUAD, UCEC, and LUSC

We extended the construction of T-GDN to four additional tumor localizations: HNSC, LUAD, UCEC, and LUSC, using TCGA data [16-19]. These localizations were selected based on the Supplementary Figure S4 from [9], which shows that the distance between N and T attractor centers is a characteristic feature of each type of cancer. PRAD, with closely positioned attractors, occupies the lower extreme along this distance scale. LUSC, by contrast, lies at the opposite extreme, with the farthest separation between normal and tumor expression profiles. HNSC, LUAD, and UCEC, in that order, span intermediate distances from PRAD to LUSC.

Network properties for these localizations are summarized and compared in Table I. The number of T-genes is around 6,000 in PRAD and increases with the distance between attractors, reaching 18,000 in LUSC. All networks exhibit similar sparsity. Another important characteristic is the maximum frequency of a node in the network, which ranges from 0.74 in PRAD to 1.00 in LUSC. This indicates that, in the case of well-separated attractors, perfect markers exist [9] whose sole activation determines whether the tissue is normal or tumor. In particular, *TBC1D7* and *PLSCR4* genes act as perfect markers in UCEC [9]. These genes are promising candidates for further translational investigation toward therapeutic applications.

With increasing normal-to-tumor distances, the number of isolated nodes generally decreases (from 151 and 193 in PRAD and HNSC, respectively, to 1 in LUSC) and the mean node degree increases, indicating a greater number of causal interconnections. The network diameter also increases, albeit modestly, from 2 to approximately 5 as attractor separation increases, while preserving the small-world character of each network. Overall, these features point to more structured T-GDN as the distance between the homeostatic and cancer phenotypes increases.

### 3.4 The N-Gene Deregulation Networks in PRAD and other localizations

Here, we return to PRAD as a prototypical case, where we identify 1097 N-genes [10]. To infer the links between them, and thus to map out the structure of the N-GDN, we focus on normal samples. As discussed in Ref. [10], somatic evolution of normal tissue involves only the N-genes. Indeed, the somatic evolution of a normal sample begins with a large set of active N-markers and progressively leads to their deactivation as the tissue approaches the tumor transition [10]. Therefore, our N-GDN is designed to capture this early phase of somatic evolution.

Note that a deactivation is tantamount to an expression deregulation for N-genes (see Sec. 2.1). As a consequence, the deregulation and activation frequencies are complementary (summing to 1); denoting the activation frequency as v_N_, the deregulation frequency is 1 - v_N_. Since activation frequency is what is bounded from below in the selection of N-genes, we naturally have an upper limit for deregulation frequencies equal to 0.95. Despite the fact that the minimum deregulation frequency for N-genes is unconstrained, we find its value to be quite large, i.e., 0.69. Since the initial fraction of active N-genes during somatic evolution is not directly observable, it is difficult to determine whether this minimum frequency reflects an advanced stage of the normal samples in our dataset or heterogeneous N-activation patterns across samples at early stages of somatic evolution. The minimum frequency takes place for *SEPTIN10*, which is an NT-gene.

A notable feature in the N-GDN is the presence of block genes [10]—i.e., genes with identical expression profiles across normal samples, indistinguishable from one another. For each block of genes sharing a specific deregulation profile across normal samples, we include a single representative node in the N-GDN. As a result, there are only 499 nodes in our network, with the largest block-node encoding over 200 genes [10]. This may be related to limited normal sample numbers, but we prefer to ascribe compressed nodes to the saltatory nature of somatic evolution reflected in multi-step cancer models [22], where expression rearrangements across many genes may occur in a coordinated manner [10].

Following the causal discovery procedure, we obtain an N-GDN, which is a directed acyclic graph. A single node remains isolated, likely due to the Loevinger coefficient threshold, H_0_ = 1/2. The network contains 4984 edges, with a sparsity index of 0.0200. Additional characteristics of the N-GDN are reported in Table I and Figure 4, which highlights *SEPTIN10* as the gene with the lowest deactivation frequency (0.69) and an out-degree of 46.

**Figure 4:**
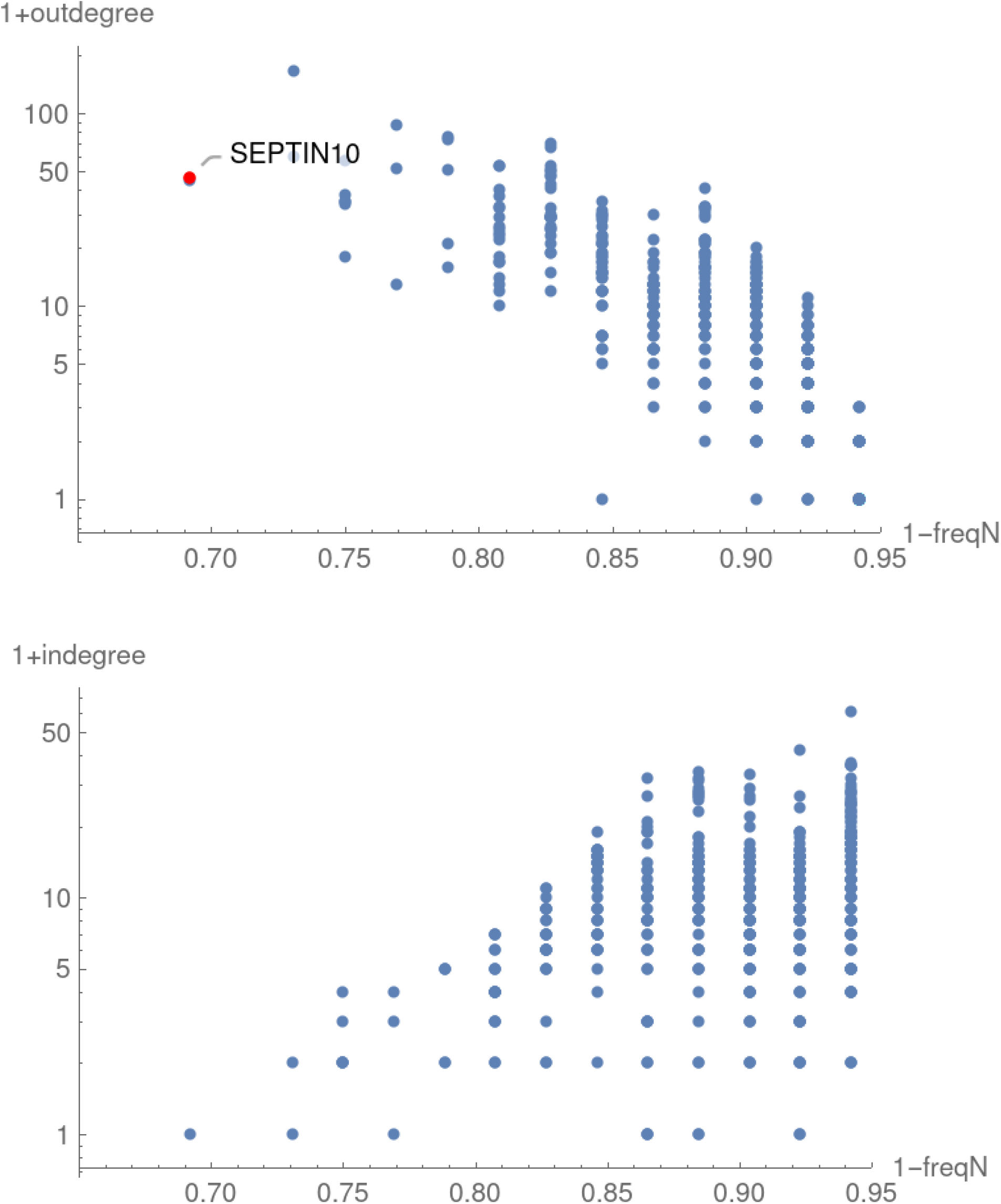
Characterization of the N-GDN in PRAD. SEPTIN10 exhibits the lowest deactivation frequency (0.69) and out-degree 46.

Comparing PRAD with other tumor localizations, we observe an important difference relative to T-GDNs: the number of N-genes remains around 1000, regardless of the distance between the attractors. This suggests that the number of N-genes might be a robust, intrinsic property of the homeostatic state of any tissue. If N-genes are interpreted as anchors of the normal state, preventing the transition of the tissue towards a tumor, then a comparable level of protection is observed for all tissue localizations. Indeed, around 500–600 N-markers are active in the least somatically evolved normal samples in the dataset irrespective of the tissue [10].

Additionally, we note that the minimum deregulation frequency of N-genes decreases as distance between attractors increase, from 0.69 in PRAD to 0.1 in LUSC. The case of UCEC, with an observed minimum frequency of exactly 0, may be a consequence of the low number of normal samples in this case (only 23).

### 3.5 Still Active Markers in Normal Samples

A distinctive feature of somatic evolution is that the deactivation of N-genes removes information on previously active genes. Whereas tumor samples reveal visible deregulation cascades, normal samples only reflect currently active nodes and, if mapped onto the N-GDN, the potential effects that would arise if these nodes were deactivated.

This is illustrated in Figure 3 (bottom panel), showing the normal sample paired with the tumor sample from the top panel. Only 34 N-genes remain active in this sample, indicating an advanced state of somatic evolution [10]. Two NT-genes common with the tumor sample are highlighted in red.

Note that on active nodes there is also pressure exerted by previously deactivated nodes. Indeed, the bottom just as the top panel of Figure 3, are both partial representations of their respective GDN. Influence from already deactivated nodes within the N-GDN is difficult to illustrate, precisely because all records of the originally active nodes are erased in the course of somatic evolution.

### 3.6 Evolution Dynamics Dictated by the GDNs

The possibility of reversing the tumor phenotype by reactivating the N-gene network through targeted interventions raises a fundamental question about the asymmetry of the underlying dynamics. Such a process is not spontaneous: external perturbations are required to activate a small subset of N-genes, which may then trigger a widespread reactivation of the network. Why, then, is this asymmetry present? Why can an active N-gene become deactivated spontaneously, whereas a deactivated N-gene does not reactivate under the same conditions? What drives the effectively irreversible progression of normal samples toward N-marker loss and ultimately tumor formation?

Small irreversible changes—such as mutations or stable epigenetic alterations—provide a plausible mechanism. By enforcing the deactivation of N-genes, the accumulation of these perturbations over the course of somatic evolution introduces a normal-to-tumor directionality into the dynamics, preventing spontaneous reversibility. Additionally, tumors exhibit higher entropy than normal tissues [23,24], further biasing the evolution toward the tumor state.

The data-inferred N- and T-GDNs, in that order, describe the spontaneous processes of N-gene deactivation and T-gene activation, respectively [10]. We assume that processes involving targeted interventions, such as gene knockdown or forced expression of genes, which go against spontaneous processes, are described by the reversed networks. For example, a forced deactivation of a T-gene may act on its parent genes defined by the T-GDN. The argument supporting this assumption is that the Loevinger coefficients for the direct and reverse processes are equal.

Modeling the dynamics of somatic evolution, or those ensuing from gene interventions, in a very simplified setting can proceed as follows: (1) start with an initial configuration of activated genes (e.g., derived from a sample); (2) establish a fictitious time axis related to the Monte Carlo steps, anologous to Glauber dynamics [25]; (3) stochastically evolve the system by allowing activation and/or deactivation events according to direct and reverse networks.

Such a dynamics is completely constrained by the initial state and structure of the networks. In particular, we neglect spontaneous activations or deactivations, which are nevetheless important, as shown in Fig. 2. The resulting simulation scheme is much simpler than, for example, a Gillespie algorithm for reactions [26].

In the dynamic simulation, the following set of rules governing spontaneous and forced transitions in the N- and T-gene systems hold:

Spontaenous dynamics.

- **r1**. In normal samples, the spontaneous process is the deactivation of N-genes, and deactivation cascades propagate in the direction of decreasing v_N_.
- **r2**. In tumor samples, the spontaneous process is the activation of T-genes, and activation cascades propagate in the direction of increasing v_T_. Forced dynamics consistent with spontaneous directionality.
- **r3**. Forced processes of the same character as spontaneous ones—i.e., a deactivation of an active N-gene or an activation of an inactive T-gene—follow the same direction of their spontaneous analog. Rates may differ, but we are not currently in the position of determining particular rate values. Forced dynamics opposing spontaneous directionality.
- **r4**. Forced processes of opposite character to spontaneous ones generate cascades in the reverse direction. The activation cascade in the N-GDN propagates toward increasing v_N_, while the deactivation cascade in the T-GDN propagates toward decreasing v_T_. Cascade propagation rules.
- **r5**. Forced activation of an N-gene in a normal sample triggers an activation cascade involving N-genes of activation frequencies higher than the initial gene.
- **r6**. Forced activation of a T-gene in a normal sample triggers an activation cascade involving T-genes of activation frequencies higher than the initial gene.
- **r7**. Forced T-activation of an NT-gene in a normal sample triggers cascades of N-gene deactivations and T-gene activations with the appropriate frequencies.
- **r8**. Forced activation of an N-gene in a tumor triggers an activation cascade involving N-genes of activation frequencies higher than the initial gene.
- **r9**. Forced deactivation of a T-gene in a tumor triggers a deactivation cascade involving T-genes of activation frequencies lower than the original.
- **r10**. Forced N-activation of an NT-gene in a tumor triggers cascades of N-gene activations and T-gene deactivations with the appropriate frequencies.

As an illustrative example, let us simulate the knockdown of *EPHA10* in a PRAD tumor sample. *EPHA10* is the T-gene with the highest activation frequency in the PRAD dataset and the first, from a panel of 8, we used in Ref. [10] to classify PRAD tumors. In particular, it belongs to the tumor-above category, therefore, a knockdown deactivates this gene, consistently with its known oncogenic role [27]. According to rule **r9**, a deactivation cascade advancing toward lower T-activation frequencies should follow the knockdown of *EPHA10*.

In this example, we initialize the system with the deregulation profile of a specific tumor sample from our dataset, comprising 2640 active T-genes, including *EPHA10*. We then enforce the permanent deactivation of *EPHA10*. In addition to the reverse dynamics introduced by the intervention on *EPHA10* (following **r9**), we let the direct process to act at a rate 5 times slower to simulate the response of the tumor to the intervention (following **r2**).

With this configuration, we perform 500 simulations to estimate the average activation state of all T-genes as a function of time (number of iterations). The results are shown in Figure 5, top panels. After 100 iterations, we observe a reduction in the average activation of active genes, reflecting the effect of the intervention. However, after 1000 iterations, the average activation increases for many initially inactive T-genes, indicating tumor recovery and escape.

**Figure 5:**
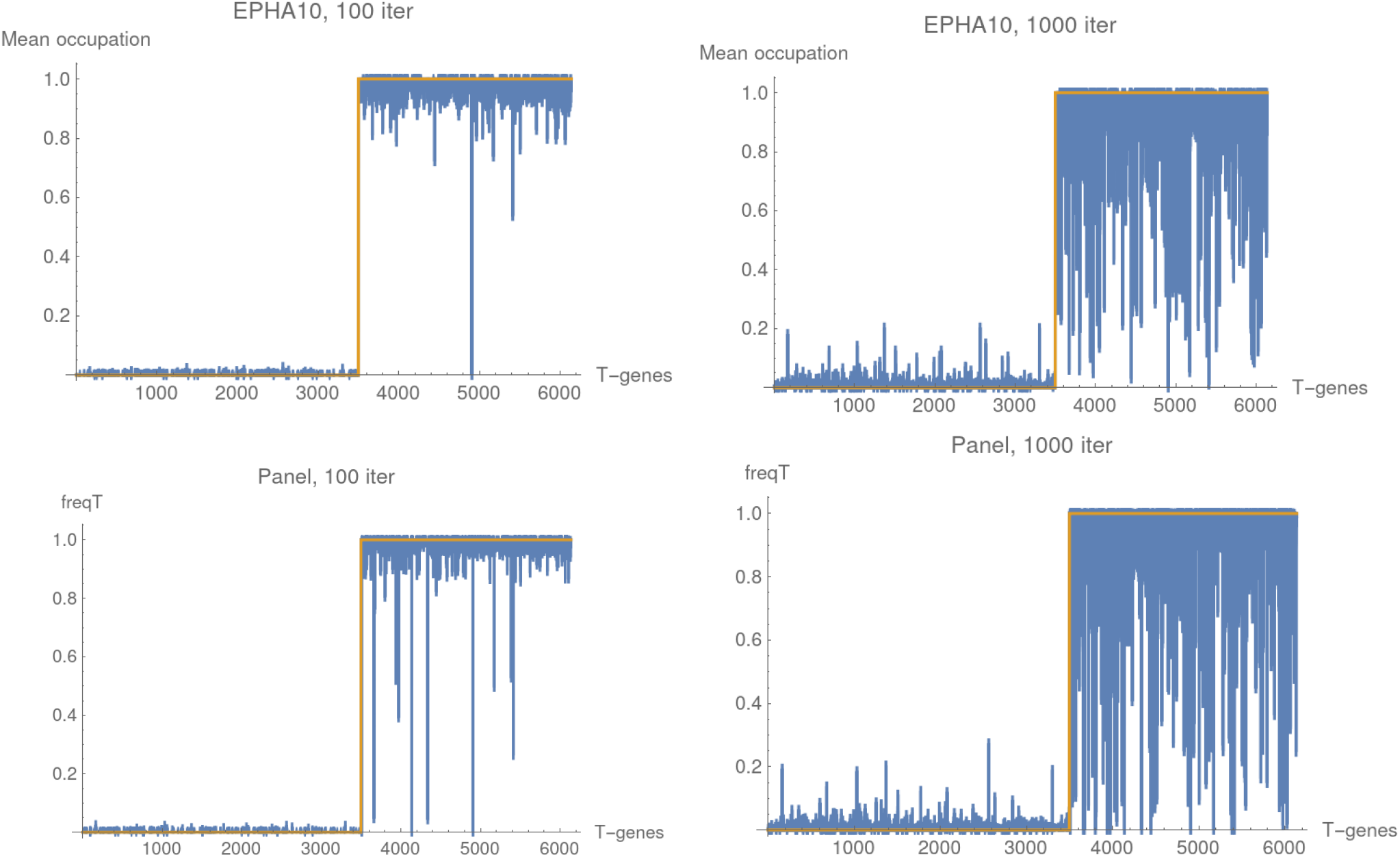
Simulated evolution dynamics in PRAD. **Top panel**: Evolution of a tumor sample after forced deactivation of the EPHA10 gene. The number of iterations, 100 and 1000, play the role of time. **Bottom panel**: Evolution of the tumor sample when the complete panel of 8 genes is forcedly deactivated. Escape is also apparent after 1000 iterations.

In the bottom panels of **Figure 5**, we consider a second example. The same sample is used as initial condition, but in this case the deactivation of the whole panel of 8 T-genes is enforced. Because the panel is perfect, that is at least one gene is active in every tumor sample, one would naively expect that targeting the full panel would eliminate the tumor; however, this is not the observed situation. After 1000 iterations, the system exhibits again tumor recovery and escape.

The failure of the intervention can be explained by the structure of the T-GDN. Indeed, at least one gene in active in every tumor sample, but this does not imply that the reverse cascade which starts from the panel genes may reach the whole T-GDN. The portion of the network that is not reached provides an escape route for the tumor.

As a third example, we check the dynamics dictated by our T-GDN against an experiment on knockdown of the transcriptional regulator *POM121* gene on two prostate cancer lines [28].

In the experiment, each cell line includes four control and four replicates in which *POM121* is knocked down. This sample size is limited and does not allow to define characteristic expression intervals for T-genes. Therefore, we rely on the T-gene identification based on TCGA data and assess changes in gene activation by comparing log-fold differential expression values in the experiment (cancer cell replicates relative to control cells) with those observed in TCGA data (tumor samples relative to normal samples). When the measured log-fold differential expression exceeds the corresponding TCGA reference (in absolute value), we register a change in the gene activation. For example, for a tumor-above gene, a negative log-fold differential expression whose absolute value surpasses the TCGA reference is interpreted as a deactivation. This analysis is intended as a qualitative consistency check; a more extensive validation of our GDN framework will be required.

Paired histograms of *POM121* expression in normal and tumor samples of TCGA-PRAD data are shown in the top panel of Figure 6. These distributions indicate that *POM121* behaves as a tumor-outside gene, however with insufficient samples in the lower interval. Accordingly, it is classified here as a tumor-above T-gene with a frequency 0.248, intermediate between the minimum (0.1) and the maximum value (0.74) exhibited by the *EPHA10* gene. The mean log-fold differential expression is 0.258.

**Figure 6:**
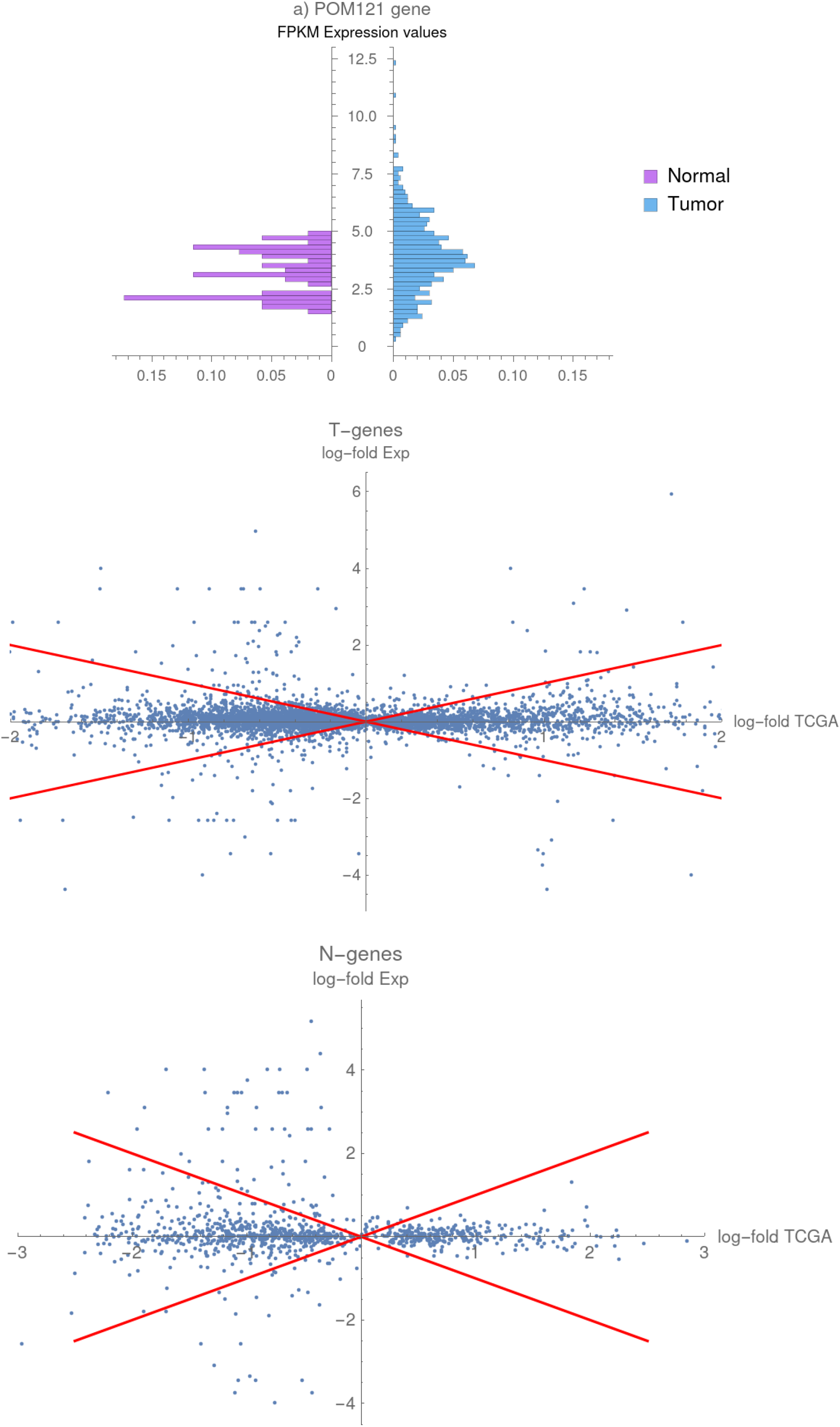
Qualitative check of the predictions of T-GDN against the experiment of POM121 knockdown in PRAD cell lines. **Top panel**: Paired histograms for the POM121 gene according to the PRAD-TCGA data. **Center panel**: Comparison between log-fold change of expressions for T-genes in the experiment and in the TCGA data for the X22RV1 cell line. This comparison is used to define activated or deactivated genes. **Bottom panel**: Comparison between log-fold change of expressions for N-genes in the experiment and in the TCGA data for the X22RV1 cell line.

We find that 5520 of the TCGA T-genes and 1009 of the TCGA N-genes are measured in the experiment. For T-genes, we distinguish activation and deactivation events taking place for genes of greater or lower activation frequency than that of *POM121*’s. According to rule **r9**, deactivations are expected to preferentially affect genes with frequencies lower than that of the intervened gene. For N-genes, in contrast, no active genes are available for deactivation, and no newly activated genes are expected (they are possible only if the intervened gene has NT-like character, according to rule **r10)**.

Figure 6, center panel, shows a plot of log-fold differential expression for the T-genes in the X22RV1 cell line. The bottom panel is the similar picture for N-genes. The number of activated and deactivated genes for both experimentally studied cell lines is given in Table II. These numbers are normalized to the total number of genes in each class (T-genes lower frequencies, T-genes higher frequencies and N-genes). Random events would lead to similar fractions in all classes.

**Table II:**
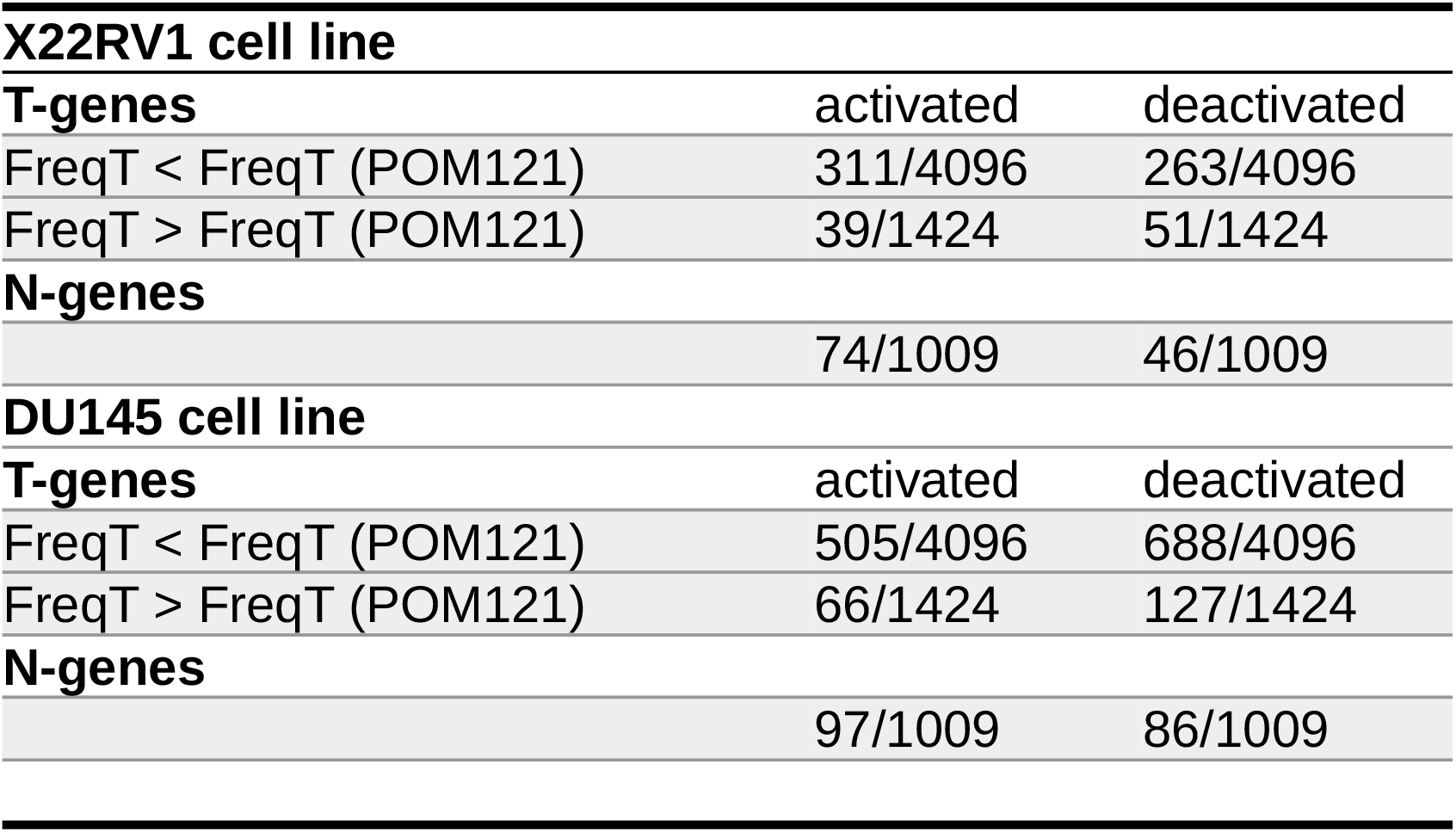
Qualitative check of the predictions of T-GDN against the experiment on the knockdown of the POM121 gene in two PRAD cell lines. The number of T- and N-gene activations or deactivations is shown.

Note that T-gene deactivations for frequencies below *POM121*’s frequency are indeed favored. A chi-squared test yields a p-value on the order of 10^{-14}, indicating a significant association between frequency intervals and activation/deactivation patterns.

The second point to notice is that 46 N-genes are deactivated. Under the assumption that no N-genes are active in a tumor sample, the fraction of deactivated N-genes can be interpreted as an estimate of the noise level induced by the limited sample size and the non-canonical criterion used for determining changes in gene activation. With regard to activations, the interesting point is the observation of N-gene activations. Their number is small, 74, but their fraction is 1.6 times the noise level. We can not decide whether it is simply noise or a true violation of the **r10** rule. In the later case, activation events in the N network could be introduced through the deactivated NT-genes in the T network.

Finally, we examined how many genes in the inverse T-GDN cascade are contained among the registered deactivations of T-genes below the reference frequency. We identify 18 such genes, all of which exhibit a causal connection with *POM121*: {*GOSR1, MTRR, MAML1, NCOA6, NUP155, ZNF792, ZNF33A, EMSY, GPATCH2, KIAA0100, VEZT, MPHOSPH9, VPS13A, SOGA1, POLR3B, PRKAA1, NBN, BACH1*}.

In general, normal and tumor evolution dynamics are the simplest processes. A more challenging task is the modeling of the normal-to-tumor transition, involving both (direct and inverse) N and T networks. NT-genes would serve as connecting elements. Spontaneous gene activations might be included. This same framework should apply to modeling partial tumor phenotype reversal [26], where forced activation or deactivation of one or more markers can induce partial reversion. AGER represents a paradigmatic experimental system [30].

## 4. Overview & Concluding Remarks

The Gene Deregulation Network framework introduced here offers several advantages over traditional gene regulatory network approaches. First, by focusing on deregulation rather than functional relationships, it bypasses the limitations imposed by incomplete gene annotation. Second, the separation into N and T networks based on tissue markers provides a natural decomposition aligned with the biological distinction between somatic evolution in normal tissues and clonal evolution in tumors.

The causal inference approach, grounded in the probabilistic theory of causation and employing the Loevinger coefficient, yields directed acyclic graphs that capture asymmetric relationships between genes. The sparsity of these networks (0.27% of possible edges in T-GDN, 2.0% in N-GDN) indicates, that despite the large number of genes involved, deregulation follows structured pathways rather than random connections.

Several observations merit further investigation. The high out-degree of low-frequency genes suggests that rarely deregulated genes may function as “master regulators” capable of influencing multiple downstream targets. Conversely, the high in-degree of high-frequency genes highlights the presence of hubs where multiple deregulation pathways intersect—potentially marking core tumor dependencies.

The presence of block nodes in the N-network, particularly the large cluster containing over 200 genes with identical expression profiles, raises interesting questions about the nature of somatic evolution. This may reflect coordinated regulation or, alternatively, suggest that normal samples feature discrete transcriptional profiles corresponding to progressive stages of somatic evolution.

Analysis of tumor samples reveals a transition from predominantly spontaneous activation in early tumors to cascade-driven activation in advanced disease. This finding has implications for understanding tumor progression and potentially for therapeutic timing—interventions targeting cascade mechanisms may be more effective in advanced tumors, while early tumors might require different strategies.

The reversibility associated with the Loevinger coefficient between direct and inverse processes suggests that the underlying regulatory logic is symmetric, with directionality imposed by external factors such as mutations. Combined with the entropy difference between normal and tumor states, this supports a thermodynamic perspective on cancer progression that warrants further theoretical development.

Finally, the potential for modeling partial phenotype reversal through forced marker activation or deactivation opens avenues for the rational design of differentiation therapy approaches. The *AGER* experiment [30] provides a paradigmatic example of how single-marker perturbations can propagate through the network to induce broader phenotypic effects.

## Acknowledgements

Authors are grateful to R. Perez for a critical revision of the manuscript. The authors acknowledge support by the Financial and International Projects Office of the Ministry of Sciences, Cuba (project PN692LH007-095).

## Author Contributions

F.C. and R.H. computed the networks, contributed to network analysis and visualization and manage the GitHub repository. J.P.G. developed the causal discovery code. G.G. ideated the causal discovery scheme and introduced the basic concepts of N- and T-genes. A.G. designed the research plan and wrote the initial draft. All authors analyzed and interpreted the results, contributed to the manuscript, and approved the final version.

## Data Availability

All data used in this study are publicly available through the TCGA Research Network (https://www.cancer.gov/tcga). Code and processed network files are available at https://github.com/[repository] upon publication.

